# Pam3CSK4 as a Cross-Species Adjuvant for Polysaccharide Vaccines: Efficacy in Humanized Mouse and Non-Human Primate Models

**DOI:** 10.1101/2025.10.09.681501

**Authors:** Jamie E. Jennings-Gee, Alexis E. Adams-Sims, Karen M. Haas

## Abstract

Polysaccharide-based vaccines are critical for preventing bacterial infections, yet their efficacy is often limited by weak antibody responses. Unfortunately, efficacious adjuvants for licensed native polysaccharide vaccines are lacking. The TLR4 agonist, monophosphoryl lipid A (MPL), significantly increases antibody responses to capsular polysaccharides in mice via B cell-intrinsic TLR4 and MyD88-dependent signaling. However, due to the lack of TLR4-driven adjuvant effects on polysaccharide-specific responses in non-human primates and the limited responsiveness of human B cells to TLR4 agonists, we sought to identify alternative MyD88-activating TLR agonists that could serve as suitable adjuvants to enhance humoral responses to polysaccharide vaccines in humans. In vitro assays revealed the TLR1/2 agonist Pam3CSK4 synergized with strong BCR crosslinking to optimally enhance both mouse and human B cell activation and antibody secretion. In vivo, Pam3CSK4 alone had no effect, but when paired with a squalene-based emulsion significantly increased polysaccharide-specific antibodies in both immunocompetent and PBMC-humanized mice that proved highly protective against lethal pneumococcal infections. Although a dual TLR2-7 agonist showed similar potent in vitro activity, it failed to enhance polysaccharide-specific IgG responses in vivo, mirroring the antagonistic effects observed when TLR2 and TLR7 agonists were combined both in vitro and in vivo. By contrast, inclusion of Pam3CSK4 in an adjuvant containing MPL, synthetic cord factor, and squalene emulsion further augmented protective polysaccharide-specific antibody responses in mice and rescued adjuvant effects in non-human primates. These findings reveal Pam3CSK4-containing formulations as promising adjuvants for native polysaccharide vaccines, with strong translational potential to enhance humoral immunity in humans.

**One Sentence Summary:** Pam3CSK4-based adjuvants show promise for boosting protective antibody responses to polysaccharide vaccines.

## INTRODUCTION

Antibody (Ab) responses to bacterial capsules and other pathogen-associated polysaccharides are essential for host protection. Polysaccharide antigens (Ags) typically behave as T cell-independent type 2 (TI-2) Ags, which elicit rapid Ab responses primarily through innate-like B cell subsets, including marginal zone and B1 B cells (1–9). While TI-2 Ags can induce Ab (predominantly IgM) responses independently of cognate T cell help, T cells may still contribute to class switching and IgG production through bystander activation and/or non-classical T cell help (10, 11). Given the superior effector functions of IgG—including FcγR-mediated phagocytosis and complement synergy (12–14)— enhancing IgG responses to polysaccharide vaccines remains a key goal.

Current polysaccharide vaccines are typically administered as native Ags without adjuvants or as protein-polysaccharide conjugates with adjuvants. Despite their success, conjugate vaccines are costly and complex to manufacture and distribute. Adjuvants that can boost IgG responses to native polysaccharides offer a promising alternative for cost-effective coverage and rapid deployment. Traditional adjuvants like alum fail to enhance primary TI-2 responses (15), and co-administration of TLR agonists in mice is ineffective unless administration is delayed by several days (16–18), which is impractical for clinical implementation. Clinical trials with IL-12, GM-CSF, QS-21, CpG, and others have similarly failed to improve IgG responses to native pneumococcal polysaccharides (PPS) (19). Early work with Ribi, consisting of *Salmonella typhimurium* monophosphoryl lipid A (MPL) and mycobacterial cord factor (trehalose-6,6’-dimycolate) in squalene oil-in-water emulsion increased primary PPS-specific Ab responses in mice (20, 21). We previously demonstrated a related, but less toxic adjuvant consisting of *Salmonella minnesota* MPL, synthetic cord factor analog (synthetic trehalose dicorynomycolate; sTDCM, or D-(+)-trehalose-6,6’-dibehenate; TDB), and squalene emulsion significantly augments Ab responses to native polysaccharides in mice, as demonstrated by significantly increased IgG production, memory formation, and Ab secretion following boosting (4, 22–24). Using a mouse model, we identified a critical role for MyD88-driven signaling on B cells for these effects (22), and determined that B-1b and marginal zone B cells were major contributors to the increased Ab produced (4). Finally, we showed iNKT cells were critical for the adjuvant effects on increasing polysaccharide-specific IgG in mice (24).

Despite its potency in mice, in the current study, we found the MPL-TDCM-squalene emulsion adjuvant failed to promote polysaccharide-specific Ab responses in non-human primates. Our analysis revealed MPL is responsible for the majority of the adjuvant’s effect in mice. This provided the rationale for studies aimed at identifying alternative MyD88-activating TLR agonists to co-stimulate polysaccharide Ag-activated human B cell Ab production. Herein, we describe the potential for distinct TLR agonists to activate as well as antagonize polysaccharide-specific B cell responses and identify Pam3CSK4 as a TLR agonist that significantly promotes polysaccharide-specific Ab production by mouse, non-human primate, and human B cells. These findings may be leveraged to develop strategies to improve polysaccharide vaccine responses in humans.

## RESULTS

### MPL combined with squalene emulsion significantly increases Ab responses to PPS and other TI-2 Ags in mice and depends on B cell intrinsic TLR4-MyD88 signaling

Our previous work demonstrated that a combination of MPL, TDCM (or TDB), and squalene emulsion significantly increases primary and secondary Ab responses to pneumococcal polysaccharide (PPS) and other TI-2 Ags, including haptenated Ficoll. This effect was strongly dependent on TLR4 and B cell-expressed MyD88, with limited dependency on Trif signaling (22). Given the strong dependency on TLR4 and B cell-expressed MyD88, we investigated the extent to which MPL + squalene emulsion alone could function as an adjuvant for TI-2 Ab responses. We found MPL + squalene emulsion significantly increased PPS3-specific IgM and IgG responses to Pneumovax23 (Fig. 1A) and significantly increased protection against intranasal challenge with 10^7^ CFU of serotype 3 *Streptococcus pneumoniae* (WU2) (Fig. 1B) in a manner similar to MPL+ TDB + squalene emulsion. Consistent with our previous findings for MPL +TDCM + squalene emulsion (4, 22, 24), MPL + squalene-induced increases in PPS3-specific IgG levels induced by depended on B cell-expressed MyD88 as evidenced by the lack of adjuvant effect in bone marrow chimera mice reconstituted with MyD88-deficient B cells relative to chimeras reconstituted with WT B cells (Fig. 1C), although PPS3-specific IgM responses were nonetheless increased during primary responses in MyD88-deficient B cell chimeras but limited boosting was observed relative to WT chimeras. Notably, MPL + squalene significantly increased IgM and IgG responses to NP-Ficoll in B cell-deficient mumt mice reconstituted with B cells from WT and Trif^-/-^ mice, but not TLR4^-/-^ mice (Fig. 1D). Taken together, these data support that MPL combined with squalene functions as an adjuvant for TI-2 Ab responses, with effects dependent on B cell-expressed TLR4 and MyD88.

**Figure 1.**
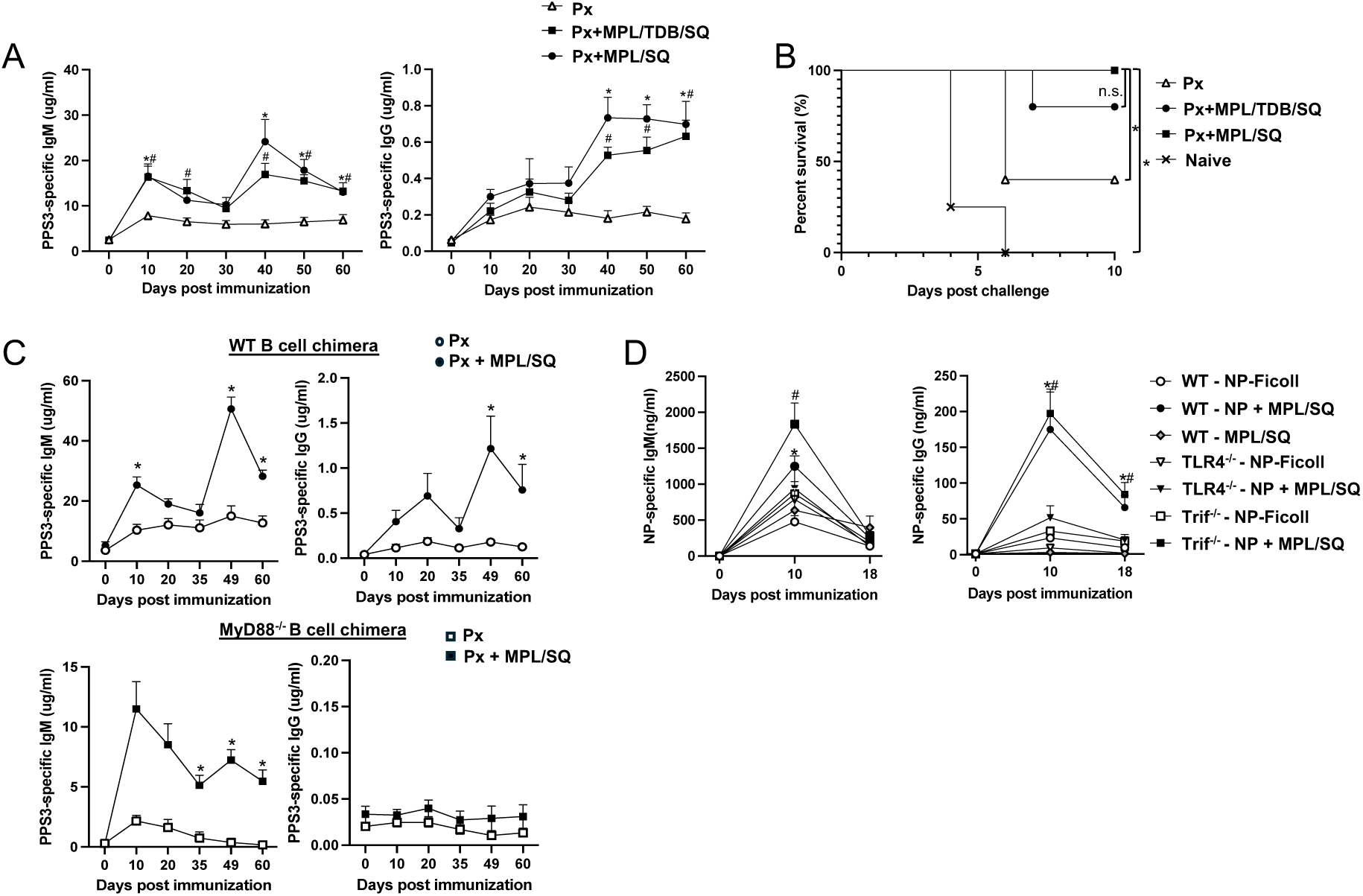
MPL + squalene emulsion functions as an adjuvant for polysaccharide-specific Ab responses and requires TLR4-MyD88 signaling on B cells. **A)** PPS3-specific IgM and IgG levels in C57BL/6 wild type (WT) mice immunized with Pneumovax23 (Px) alone or combined with MPL + squalene emulsion (SQ), or MPL + TDB + SQ i.m., with a boost on d30 (n= 5 mice/group). Analysis by mixed effects model with Bonferroni’s post-hoc analysis (*indicates difference between Px and Px with MPL/TDB/SQ and #indicates differences between Px and MPL/SQ induced Ab levels; n=5 mice/group). **B)** Survival following intranasal challenge with 10^7^ CFU serotype 3 *S. pneumoniae* (n=5 mice/group; Log rank analysis results indicated, p<0.05). Results representative of 2 independent experiments. **C)** PPS3-specific IgM and IgG levels in bone marrow chimeras reconstituted with WT:mumt bone marrow or MyD88: mumt bone marrow (20:80). *Indicates significant difference based on repeated measures ANOVA with Bonferroni’s post-hoc analysis (n=8 WT B cell chimeras and 4 MyD88 B cell chimeras per group). **D)** NP-specific IgM and IgG responses in mumt mice reconstituted with 3×10^7^ WT, TLR4^-/-^, or Trif^-/-^ B cells i.v. and immunized with 5 μg NP-Ficoll i.m. alone or mixed with MPL+SQ. Statistical differences between groups of WT(*) or Trif-/-mice (#) that were vaccinated with Px alone versus Px+adjuvant as assessed by repeated measures ANOVA and Bonferroni’s post-hoc analysis are indicated (n=7 mice/group).

### The MPL-based adjuvant augments PPS Ab responses in mice but not NHP

Immunization of adult female African green monkeys (AGM) with Pneumovax23 alone or with MPL+TDCM+squalene emulsion adjuvant did not yield significant differences in whole Pneumovax23- or PPS3-specific IgM, IgG, or IgA levels between weeks 1 and 5 post immunization (Fig. 2A-B). We failed to find increases in Ab responses to PPS-4, −14, −19F, or −23F (Supplemental Fig. 1). The AGM used were 17-20 years old, which may be considered the equivalent of 50-65 year old humans (25). We think it is unlikely the lack of effect in NHP was attributable to advanced age, as the adjuvant significantly increased Pneumovax23- and PPS3-specific IgM levels during primary responses in 20-24 month old female C57BL/6 mice, which approximate 60-70 year old humans (Fig. 2C-D). Significant increases in Pneumovax- and PPS3-specific IgG were also observed in aged mice following boosting. Thus, the MPL-squalene emulsion adjuvant significantly increased protective pneumococcal capsule-specific Abs in mice, including aged mice, through a mechanism requiring TLR4-MyD88 signaling on B cells, but did not significantly increase polysaccharide-specific Ab responses in NHP.

**Figure 2.**
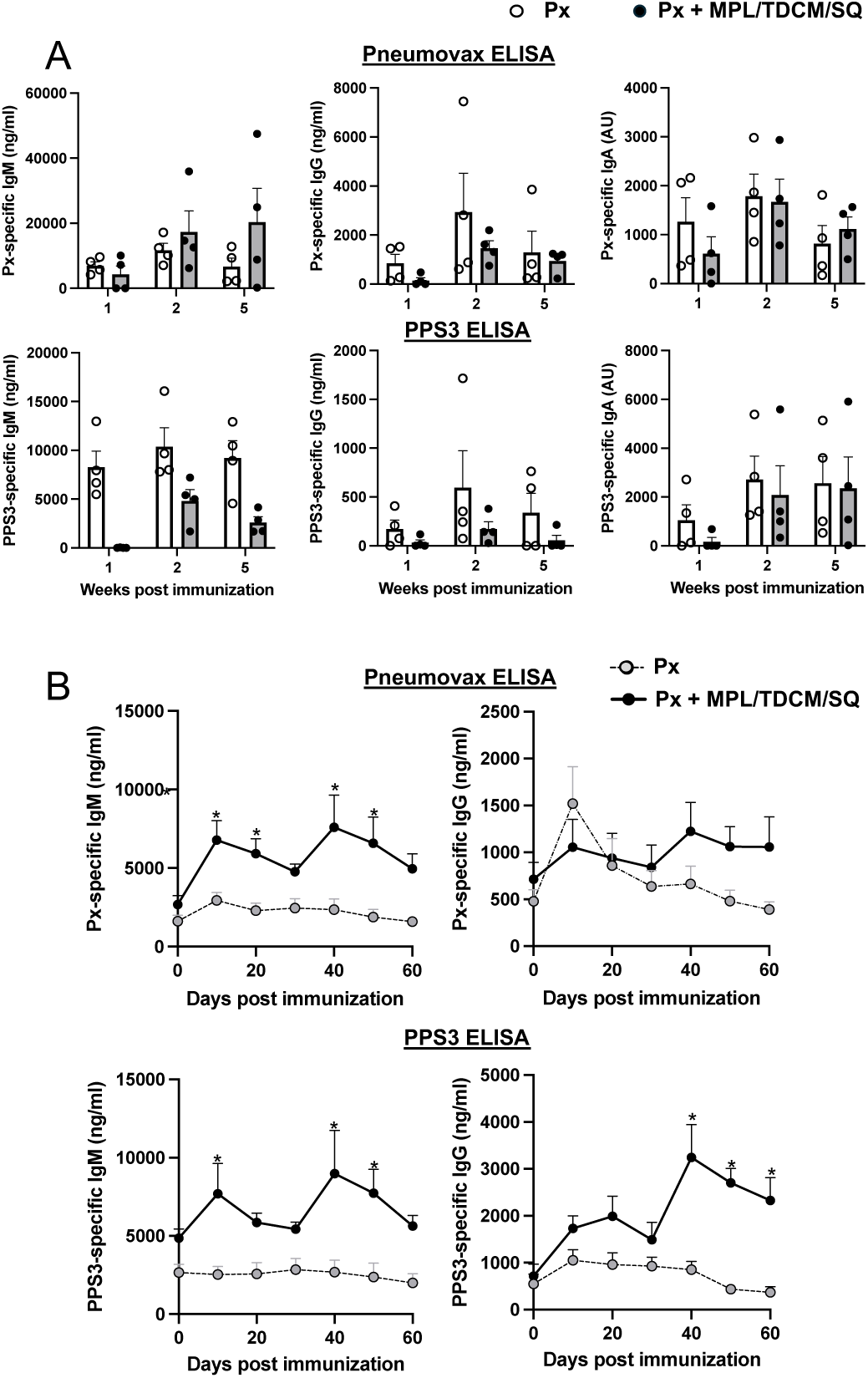
MPL+ TDCM + squalene emulsion does not increase PPS-specific Ab responses in mature African Green monkeys but does so in old mice. **A)** Whole Pneumovax (Px)- and PPS3-specific IgM, IgG, and IgA responses in 17-20 year old female AGM at 1, 2, and 5 weeks post-vaccination (i.m.) with Pneumovax23 containing 12.5 μg each PPS alone or mixed with MPL+TDCM+squalene emulsion (SQ). Individual baseline Ag-specific Ab values were subtracted from week 1, 2, and 5 Ab values to determine vaccine-induced increases in PPS-specific IgM, IgG, and IgA levels. **B)** Px- and PPS3-specific IgM and IgG responses in 20-24 month-old female C57BL/6 mice immunized (i.m.) with Pneumovax23 containing 0.125 μg each PPS alone or mixed with MPL+TDCM+SQ on d0 and 30. Differences between groups are indicated by asterisks (repeated measures ANOVA with post-hoc Bonferroni’s analysis, p<0.05, n=6 mice/group).

### Evaluation of TLR agonists for co-activating human B cells in the context of BCR crosslinking

Given the requirement for B cell-specific TLR4 expression in supporting MPL-based adjuvant effects, the lack of effect in NHP, and the negligible expression of TLR4 on human versus mouse B cells (26, 27), we sought to identify other MyD88-activating TLR agonists that would potentiate increased TI-2 Ag-specific B cell activation and thereby promote increased primary and secondary Ab responses to polysaccharide Ags in humans. To do this, we stimulated human PBMC and CD19^+^-purified B cells with alternative MyD88-activating TLR agonists, strong BCR crosslinking using biotinylated anti-mouse Ig (H+L) F(ab’)_2_ Abs with streptavidin, or both, and measured IgM-, IgG-, and IgA-secreting cells via ELISpot analysis 5 days later. Our goal was to identify agonists that stimulate little to moderate Ab production on their own but synergize with BCR signaling to significantly augment Ab production.

Flow cytometric analysis was performed on cultures of stimulated PBMC isolated from four donors to evaluate TLR agonist co-stimulatory potential. BCR crosslinking alone significantly increased the percentage of B cells among PBMC relative to media or streptavidin-only wells after 5 days of culture (Supplemental Fig. 2). Addition of Pam3CSK4 (TLR1/2), FSL1 (TLR2/6), and CpG (TLR9) significantly increased the percentage of CD20^+^ cells in culture relative to both TLR agonist and BCR crosslinking alone. R837 (TLR7), R848 (TLR7/8), adilipoline (TLR2/7), and flagellin (TLR5) showed a similar trend for 3 out of 4 donors whereas Addavax (squalene emulsion) had no effect. Similar trends were observed for the total number of B cells that had undergone CFSE-marked division and the overall number of IgG^+^ B cells following 5 days of culture, although only Pam3CSK4 + αIg cultures showed yields that were significantly increased over either stimuli alone across all 4 donors (Supplemental Fig. 2). Thus, although donor-to-donor variation was observed, all TLR agonists generally increased B cell numbers in PBMC cultures with BCR crosslinking, with Pam3CSK4 having the most consistent effect among donors.

Given the above results, we proceeded with ELISPOT analyses using all of the TLR agonists described above. As shown in Fig. 3A, activation of BCR signaling alone in unfractionated PBMC or purified CD19^+^ B cell cultures induced insignificant numbers of ASCs compared to media alone. Pam3CSK4 had little effect on ASC yield when used alone, but significantly increased IgM ASC numbers when combined with BCR activation in PBMC cultures, and significantly increased both IgM and IgG ASCs in purified B cell cultures, indicating a role for direct B cell stimulation in co-activation. FSL-1 induced similar increases, albeit to a lesser extent. The TLR5 and TLR7 agonists, flagellin and R837, stimulated increased IgG ASC production in the presence of BCR crosslinking, but only in PBMC cultures, suggesting these effects required agonist activity on non-B cells (28). Consistent with previous reports, when used alone, the TLR7/8 and 9 agonists, R848 and CpG, stimulated the greatest increases in IgM, IgG, and IgA ASCs in PBMC cultures relative to other agonists. Yet, with the exception of IgA ASCs, there was no further increase with these agonists when combined with BCR crosslinking. However, in purified B cell cultures, R848 and CpG synergized with BCR crosslinking to yield significant increases in IgM (R848) and IgG (R848 and CpG) ASCs. Finally, evaluation of a dual TLR2-TLR7 agonist, adilipoline, revealed its capacity to induce significant increases in IgM, IgG, and IgA ASCs in both PBMC and purified B cell cultures. Collectively, work evaluating subjects whose B cells in bulk PBMC cultures produced at least 2-fold greater ASC numbers with BCR crosslinking plus TLR agonist relative to either stimulus alone (n=10 donors) revealed adilipoline, Pam3CSK4, and R837 supported the most consistent increases among donors, with adilipoline showing the most potent effect (Fig. 3B).

**Figure 3.**
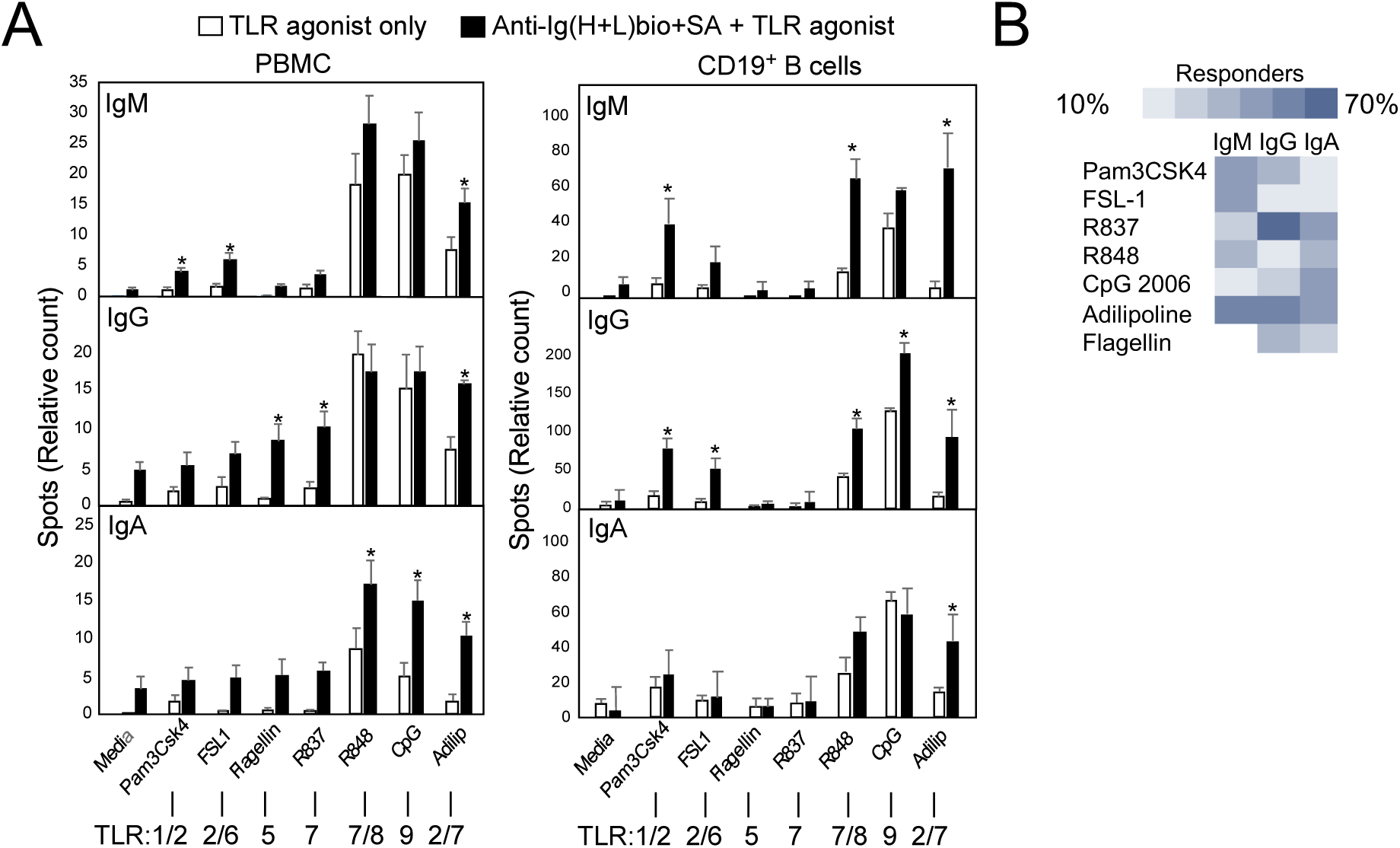
Activation of human B cell Ab secretion by TLR agonists and strong BCR crosslinking. **A)** Human PBMC (10^5^/well) or CD19-selected B cells (10^4^/well) were cultured for 5 days in rhuIL-2 (10 ng/ml)-supplemented media, Pam3CSK4, FSL1, flagellin, R837, R848, adilipoline, or CpG2006 (1-2 μg/ml) alone or in combination with 1 μg/ml biotinylated goat anti-human Ig (H+L) F(ab’)_2_ plus 5 μg/ml streptavidin. On day 5, cells were transferred into ELISPOT plates and cultured for an additional 16 hours, with total IgM, IgG, and IgA spots detected. Pooled results shown for 5 donors (3 males, 2 females). Asterisks (*p<0.05) indicate values obtained with TLR agonist plus BCR crosslinking are significantly different from values obtained for BCR crosslinking alone and TLR agonist-only stimulation. **B)** Heatmap of ASC responses of 10 donors to TLR agonists plus BCR crosslinking versus TLR agonist alone with positive responders for individual isotypes considered to have at least a two-fold increase in ASC numbers with TLR agonist plus BCR crosslinking relative to either stimulus alone.

### TLR2 and TLR7 agonists used individually, but not in combination, increase protective PPS-specific IgG levels in mice in vivo

Given the notable effects of adilipoline, Pam3CSK4, and R837 on induction of ASC in the context of BCR crosslinking on human B cells in vitro, we assessed the potential for these agonists to function as adjuvants for PPS-specific Ab responses in vivo. For these experiments, wild type C57BL/6 mice were immunized i.m. with Pneumovax23 mixed with Addavax (squalene emulsion) alone or further combined with TLR agonists (MPL, Pam3CSK4, R837, adilipoline, or Pam3CSK4 plus R837). As shown in Fig. 4A, all TLR agonists significantly increased PPS3-specific IgM levels over Pneumovax23-squalene, although MPL generally yielded higher levels than other agonists based on area under the curve (AUC) analysis. However, Pam3CSK4, R837, and MPL used alone significantly increased PPS3-specific IgG levels to at least 10-fold higher than those produced in response to Pneumovax-squalene in control animals, whereas mice immunized with the dual agonist, adilipoline or the Pam3CSK4/R837 combination failed to produce appreciable increases in PPS3-specific IgG (Fig. 4A). Similar trends were found when pooled PPSs were used in ELISA coating, although measured responses were more variable among individual mice (Supplemental Figure 3A). Flagellin similarly reduced Pam3CSK4 adjuvant-mediated increases in PPS-specific IgM and IgG levels (Supplemental Figure 3B), suggesting antagonistic effects were not limited to TLR7 agonists.

**Figure 4.**
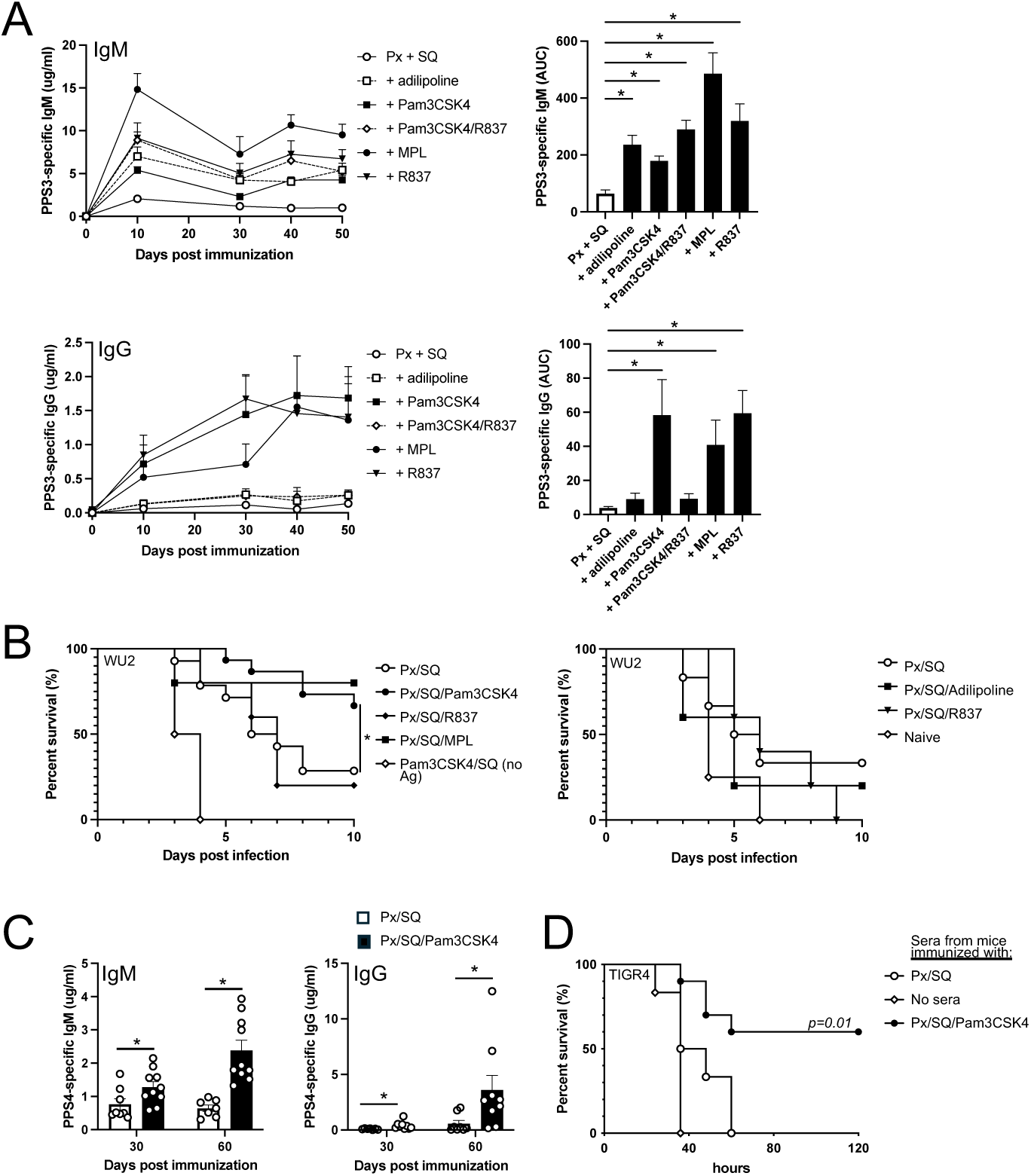
TLR2 and TLR7 agonists delivered in squalene emulsion function as adjuvants for PPS-specific Ab responses when used alone but not together in WT mice. **A)** PPS3-specific IgM and IgG responses in WT mice immunized i.m. with Pneumovax (Px) in SQ alone or combined with the indicated TLR agonists. Differences in responses between mice receiving TLR agonists versus mice immunized with Px-SQ alone were assessed by repeated measures ANOVA with Bonferroni’s post-hoc analysis. **B)** On day 75 post immunization, mice were challenged i.n. with 10^7^ CFU *S. pneumoniae* strain WU2 and monitored for survival. Results from 2 pooled replicate experiments. *Asterisk indicates Log-rank (Mantel-Cox) test significance (p-0.0187) for Px/SQ/Pam3CSK4 group (n=15) versus Px-SQ group (n=14 mice). **C)** PPS4-specific IgM and IgG responses in mice from the immunization described in (A). **D)** Survival of CD19^-/-^ mice challenged i.p. with 1000 CFU serotype 4 strain TIGR4 co-delivered with 1 μl sera from mice immunized with Px/SQ/Pam3CSK4 versus Px-SQ (n=6-11 per group). The p value indicates a significant difference in survival between Px/SQ/Pam3CSK4 versus Px-SQ immunized mice (Log rank analysis).

Interestingly, in mouse splenocyte cultures containing V_H_B1-8 Tg B cells, adilipoline, Pam3CSK4, R837 and the Pam3CSK4/R837 combination drove NP-specific IgM production when used in combination with NP-Ficoll-mediated activation (Supplemental Figure 3C). However, only Pam3CSK4 significantly increased NP-specific IgG production in NP-Ficoll-activated B cells (Supplemental Figure 3C-D); moreover, R837 significantly suppressed Pam3CSK4-mediated increases in NP-specific IgG (Supplemental Figure 3C). Similarly, Pam3CSK4-supported increases in anti-Ig-induced ASC in purified human B cell cultures were reduced when R837 was added (Supplemental Fig. 3E). Collectively, these data support TLR7 and TLR2 agonists function as adjuvants for polysaccharide Ags when used individually, but when combined, suppress the individual adjuvant effects on Ag-specific IgG responses.

In contrast to the TLR2-7 combination agonists and which did not provide increased protection against lethal respiratory pneumococcal infection with serotype 3 pneumococcus over the non-adjuvanted vaccine alone, inclusion of Pam3CSK4-squalene emulsion in the pneumococcal vaccine significantly increased protection to a level similar to that when MPL-squalene emulsion was included in the vaccine (Fig. 4B). In addition to significantly increasing PPS3-specific protective Ab responses, Pam3CSK4 functioned to significantly increase the level of protective PPS4-specific Abs as evidenced by the significantly increased protection in passive sera transfer experiments with lethal systemic serotype 4 (TIGR4) challenge (Fig. 4C-D). Importantly, PamCSK4 had no effect on PPS Ab responses if squalene emulsion was not included in the vaccination (Supplemental Fig. 3F). Collectively, this data supports the TLR1/2 agonist, Pam3CSK4, when combined with squalene emulsion, functions as a potent adjuvant to promote significantly increased production of PPS-specific Abs that provide protection against lethal pneumococcal infection relative to the non-adjuvanted vaccine in mice. However, when a TLR7 agonist is combined with Pam3CSK4, it no longer increases protective PPS-specific Ab responses.

### Pam3CSK4 plus squalene emulsion functions as an adjuvant for protective human B cell responses to pneumococcal polysaccharides

Given our in vitro results with human B cells and our in vivo results in mice, we determined whether the Pam3CSK4/squalene-based adjuvant would increase PPS-specific Ab responses in humanized NSG mice. Fresh donor PBMCs were used to reconstitute NSG mice, with recipient mice subsequently immunized with the vaccine alone, vaccine plus Pam3CSK4 + squalene emulsion, or Pam3CSK4 + squalene emulsion without antigen. Successful reconstitution was confirmed by the presence of circulating Abs (Supplemental Fig. 3G). Reconstituted mice immunized with Pneumovax + Pam3CSK4 + squalene emulsion generated significantly higher levels of total Pneumovax23- and PPS3-specific Ig by 28 days post immunization (Fig. 5A). Upon normalizing PPS-specific Ab levels to the total circulating Ab levels for individual mice to account for differences in reconstitution, we found that mice immunized with the vaccine plus Pam3CSK4 + squalene emulsion had significant increases in Pneumovax23- and PPS3-specific immunoglobulins on both days 21 and 28 post immunization (Fig. 5B). Analysis of Ag-specific IgM, IgG, and IgA indicated the majority of these Abs belonged to IgM and IgG subclasses, with the adjuvant significantly increasing total PPS- and PPS3-specific IgM responses at both d21 and d28 and for IgG responses, at d28 relative to non-adjuvanted vaccine or the adjuvant alone (Fig. 5C). IgA responses were more variable; however, the adjuvanted vaccine significantly increased IgA responses to the PPS pool on d28. Analysis of Ab responses to other individual PPS included in the vaccine revealed similar trends, with the Pam3CSK4 adjuvant significantly increasing PPS4-, PPS14-, PPS19F-, and PPS23F-specific IgM responses relative to responses to the non-adjuvanted vaccine (Fig. 5D), and IgG responses in 3 to 4 out of 5 donors for PPS14, 19F, and 23F (Fig. 5D). Importantly, consistent with the increased PPS3-specific IgM and IgG in sera of mice immunized with vaccine plus Pam3CSK4-squalene emulsion, passive transfer of pooled d21 sera from these mice (equivalent to 3 μg total Ig) yielded significantly increased protection against lethal systemic serotype 3 *S. pneumoniae* infection in immunodeficient CD19^-/-^ mice relative to sera from reconstituted mice immunized with the vaccine or adjuvant alone (Fig. 5E). Thus, Pam3CSK4 combined with squalene emulsion functions as an adjuvant that significantly increases protective PPS-specific Ab responses by human B cells.

**Figure 5.**
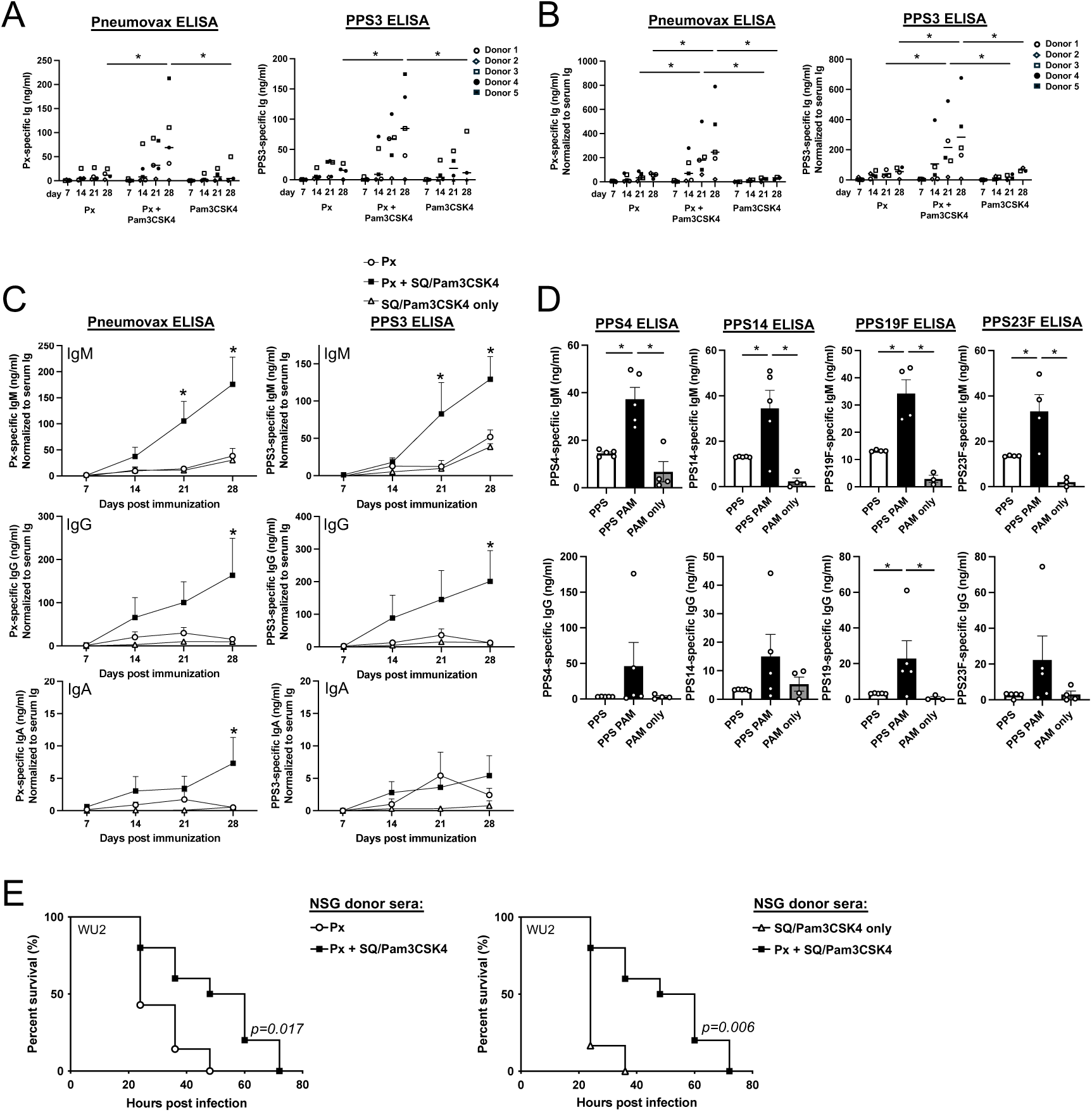
Pam3CSK4-squalene emulsion functions as an adjuvant to augment protective PPS-specific Ab responses in humanized NSG mice. NSG mice were reconstituted with human PBMCs and immunized the next day with Pneumovax (Px), Px + Pam3CSK4 + squalene emulsion (SQ), or Pam3CSK4 + SQ without antigen. A) Px- and PPS3-specific Ig levels in mice reconstituted with cells from donors 1-5. **B)** Ab concentrations in A were normalized to total serum Ig reconstitution levels for individual mice. **C)** Px- and PPS3-specific IgM, IgG, and IgA levels in NSG mice following immunization. **D)** PPS4-, 14-, 19F-, and 23F-specific IgM and IgG levels in reconstituted NSG mice on d21 post immunization, with values shown for individual donors. In A-D, asterisks indicate significant differences (p<0.05) between Px + Pam3CSK4 and Px only and adjuvant only mice (data were analyzed using mixed effects model with Dunnett’s multiple comparisons test in A-C and using a one-way ANOVA in D). Data derived from five donors, except for the Pam3CSK4 only group in which 4 donors were used. **E)** Survival following passive transfer of d21 sera (equivalent to 3 μg of Ab) from huPBMC-reconstituted immunized NSG mice into CD19^-/-^ mice infected i.p. with 100 CFU WU2 serotype 3 *S. pneumoniae*. Results are pooled from repeated experiments. Log rank analysis results are indicated (n=10 mice in Px+Pam3CSK4 group, n=7 mice in Px only group, and n=6 mice in Pam3CSK4 only group).

### Addition of Pam3CSK4 to the MPL/TDCM adjuvant increases its adjuvant effects on protective polysaccharide-specific Ab responses in mice and restores adjuvant activity in NHP

Encouraged by the positive results for Pam3CSK4 on enhancing human B cell responses to PPS in vivo, we investigated whether other agonists could be combined with Pam3CSK4 to further increase protective PPS responses in immunocompetent C57BL/6 mice. We found that TDB, similar to TDCM in our original adjuvant formulation, further increased Ab responses when combined with Pam3CSK4 and squalene emulsion (Supplemental Fig. 3H). We therefore assessed whether combining Pam3CSK4 with the commercial adjuvant containing MPL/TDCM and squalene emulsion (Sigma Adjuvant System) that was originally administered along with Pneumovax23 to the AGM monkeys with no effect (Fig. 2), improved PPS Ab responses in mice. As shown in Fig. 6, addition of Pam3CSK4 to this adjuvant significantly increased Pneumovax23- and PPS3-specific IgG responses (Fig. 6A-B). Consistent with these Ab responses, mice that received this combination exhibited the highest level of protection against high dose lethal respiratory infection with serotype 3 *S. pneumoniae* (Fig. 6C).

**Figure 6.**
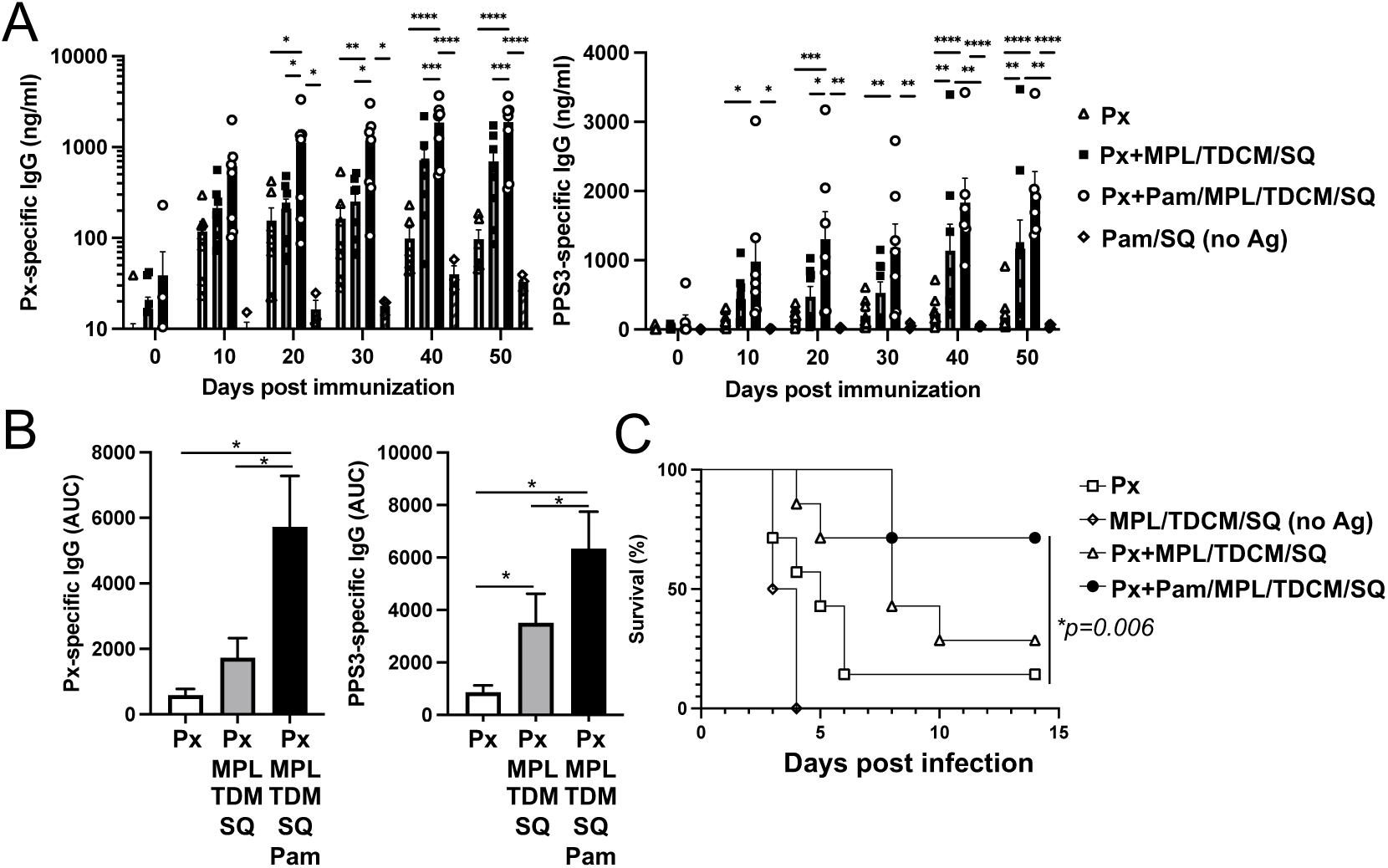
Combining Pam3CSK4 with MPL/TDCM/SQ adjuvant increases its potency for increasing PPS-specific IgG responses in WT mice. **A)** Pneumovax23 (Px)- and PPS3-specific IgG response in WT mice immunized i.m. with Px alone or Px+ MPL/TDCM/SQ, Px+ MPL/TDCM/SQ + Pam3CSK4, or Pam3CSK4 + SQ only (no antigen). All groups received a boost of same immunization on d30. *Asterisks indicate significant differences as determined by repeated measures ANOVA with post-hoc analysis (n=7 mice/group except for Pam3CSK4-SQ (no Ag) where n=4). **B)** Area under curve analysis for data in part A, with one-way ANOVA test for significance (*indicated by asterisks, p<0.05). **C)** Mice immunized as described in A were challenged i.n. with 2×10^7^ CFU *S. pneumoniae* strain WU2 on d64 post immunization and monitored for survival (*Log rank analysis with significance indicated, n=7/group).

Given that Pam3CSK4 further enhanced the response to MPL/TDCM-squalene emulsion in mice, we investigated whether inclusion of Pam3CSK4 in the MPL/TDCM-squalene emulsion adjuvant, which on its own was unable to increase PPS-specific Ab responses in AGM, would yield adjuvant effects. As shown in Fig. 7A, in contrast to results with MPL/TDCM-squalene emulsion (Fig. 2 and Supplemental Fig. 1), Pam3CSK4/MPL/TDCM-squalene emulsion significantly increased Pneumovax-specific IgM and IgA responses, as well as increased IgG responses. Although variability in responses was evident among animals, significant increases in serotype-specific Abs, including PPS4- and PPS19F-specific IgM and IgG were observed, along with significantly increased PPS3- and PPS19F-specific IgA levels (Fig. 7B-D). Collectively, the restoration of MPL/TDCM-squalene emulsion adjuvant effects upon addition of Pam3CSK4 highlights its potential to function as an adjuvant for polysaccharide-based vaccines in primates, including humans, as supported by our humanized mouse data.

**Figure 7.**
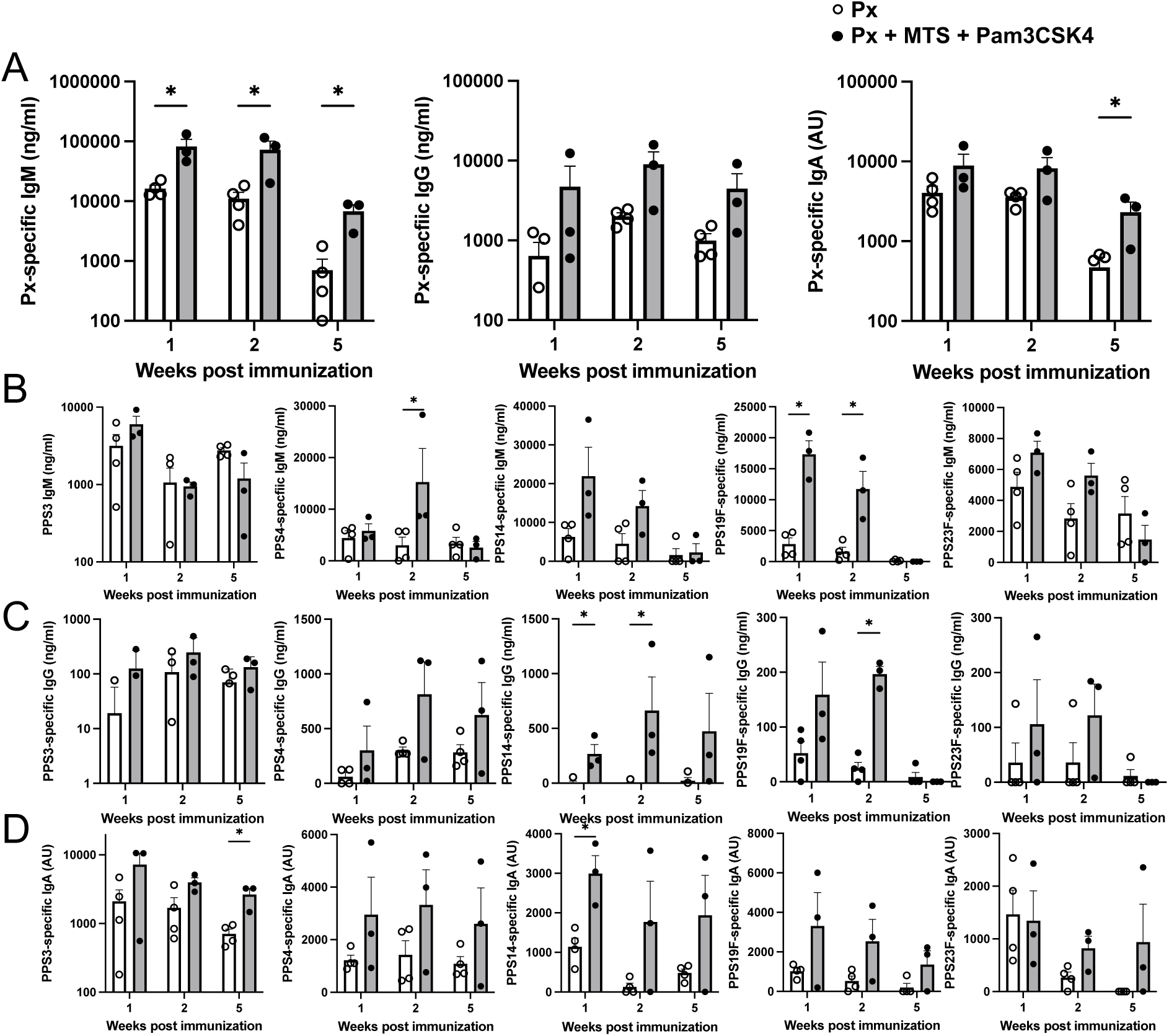
Addition of Pam3CSK4 to MPL/TDCM/SQ restores adjuvant activity for polysaccharide-specific B cell responses in NHP. **A-D)** Px-specific IgM, IgG, and IgA **(A)** and PPS3-, 4-, 19F-, and 23F-specific IgM **(B)**, IgG **(C)** and IgA **(D)** responses in 8-10 year old female AGM 1, 2, and 5 weeks post-vaccination (i.m.) with Pneumovax23 containing 12.5 ug each PPS alone or mixed with MPL/TDCM/SQ + Pam3CSK4. Asterisks(*) indicate significant differences in values as determined by unpaired t-test (p<0.05). Individual baseline Ag-specific Ab values were subtracted from week 1, 2, and 5 Ab values to determine vaccine-induced increases in PPS-specific IgM, IgG, and IgA levels. Circles represent values for individual animals.

## DISCUSSION

Polysaccharide antigens, such as those found in bacterial capsules, are critical targets for protective Ab responses, yet their T cell-independent nature—particularly in the absence of additional stimulatory signals—presents a challenge for vaccine design. TI-2 Ags primarily activate Ab production by innate-like B cell subsets, including marginal zone and B1 B cells, which rapidly produce IgM but produce modest IgG responses without additional signals (2, 9, 22, 29). Although IgM can confer protection, IgG is generally preferred in vaccine contexts due to its superior effector functions, including FcγR-mediated phagocytosis (12–14, 30). While protein-polysaccharide conjugate vaccines have successfully provided protection, especially in young children, the reliance on such complex and costly vaccines poses a potential vulnerability. The lack of suitable adjuvants for co-administration with polysaccharide vaccines represents an understudied area that could lead to development of suitable alternative routes to protection. Herein, we demonstrate a critical role for TLR4/MyD88-mediated co-stimulation of polysaccharide-specific B cell activation for a TLR4-activating adjuvant to elicit its effects in increasing protective polysaccharide-specific Ab levels in mice. Despite its potent effects in mice, we show the MPL-based adjuvant had little effect in NHP, thereby supporting the evaluation of alternative MyD88-activating TLR agonists for this purpose. Our in vitro results indicate most MyD88-activating TLR agonists have B cell stimulatory activity—either directly on human B cells or in concert with accessory cells present in PBMCs. In particular, Pam3CSK4 emerged as a potential alternative due to: 1) its ability to co-stimulate both human and mouse B cell activation, expansion, and ASC formation (including IgG production) in the presence of strong BCR crosslinking, and 2) its ability to significantly increase production of protective PPS-specific Abs in vivo in wild type mice, PBMC-humanized NSG mice, and African green monkeys. Of significant interest, we also identified antagonistic activity when either TLR7 or TLR5 agonists were combined with Pam3CSK4, whereas the opposite was true when TLR1/2 and TLR4 agonists were paired. Collectively, our findings support the strategic use of TLR1/2 agonists for next-generation adjuvant design to potentiate the protective effects of native polysaccharide vaccines in humans.

Co-stimulation of B cells through BCR and TLR provides a potent but context-dependent mechanism for modulating B cell activation, class-switch recombination, and Ab secretion. TLR and BCR signaling activates NF-kβ through distinct pathways and thus MyD88-dependent signaling may synergize with TI-2 Ag-induced BCR crosslinking to support proliferation, isotype switching, survival, and differentiation into both memory B cells and long-lived ASCs (31), especially in B-1 and MZ B cells (32), which typically contribute to TI-2 responses. In particular, previous work with mouse B cells has demonstrated BCR and TLR signaling promote enhanced CSR —for example, anti-IgD– dextran plus TLR1/2, 4, 7, or 9 ligands in the presence of IL-4 induced IgG1 CSR via NF-κB– dependent upregulation of AID (31). Although this approach stimulates extensive crosslinking, similar to TI-2 Ags with repeating epitopes, both IgM and IgD crosslinking would be expected to occur with a TI-2 Ag. Indeed, innate B cells which are largely responsible for producing Ab in response to TI-2 Ags, regardless of whether the TLR4-based adjuvant is used(4), express high levels of IgM and low levels of IgD. Solely crosslinking IgD versus both isotypes in these types of studies may skew towards follicular B cell responses and may therefore not adequately model TI-2 Ag-mediated activation of innate B cells (33, 34). Further, given the limited availability of IL-4 during these responses, CSR to IgG1 is generally limited, with IgG3, and a lesser extent, IgG2b being produced when MPL is used as an adjuvant for PPS (22). Finally, IgD engagement restrains TLR-induced plasma cell differentiation by reducing BLIMP1 expression, consistent with the suppressive effect of prolonged BCR signaling on ASC formation (31). The finding that delayed administration of TLR agonists by two days significantly increases Ab responses to PPS in mice (16–18) supports the possibility that TLR costimulatory effects on TI-2 Ab responses may be most effective after the initial phase of strong prolonged BCR crosslinking has occurred. It is therefore likely that formulations which extend TLR agonist in vivo half-life and/or potency such as squalene-based emulsions as shown in our study, or liposomes (35), contribute to the adjuvanticity of TLR4 agonists that are co-administered with polysaccharide Ags. Although work suggests physical co-engagement of the BCR and TLR synergizes to promote strong B cell activation (36, 37) and T cell-independent type 1 Ab responses when LPS or flagellin displaying multivalent epitopes are used (38), the aforementioned studies support that TLR agonists need not be linked to TI-2 Ags to function as strong adjuvants. However, our work suggests non-conventional T cells may nonetheless be required to support IgG responses (24).

Our work with MPL supports the idea that the costimulatory potential of TLR agonists for TI-2 Ab responses relies on B cell expression of TLRs. In contrast to mice whereby several TLR are expressed on naïve B cells, in humans, naïve B cells are reported to express low levels of TLR whereas activated and memory B cells express TLR1/2, 2/6, 7, and 9 (27, 39). Importantly, BCR engagement may upregulate TLR, as is the case for TLR9 (39). Indeed, human naïve B cell proliferative responsiveness to TLR1/2, 2/6, 7, and 9, but not TLR3 and 4 have been shown in the presence of soluble anti-Ig and T cell help (40). Consistent with this, along with trends for increased proliferation, we observed significantly increased Ab production in purified B cell cultures when Pam3CSK4 (TLR1/2), FSL1(2/6), R848 (7/8), CpG (9), or adilipoline (2/7) was combined with multivalent BCR crosslinking and IL-2. While our focus led us to further study Pam3CSK4 as a suitable adjuvant for human polysaccharide vaccines, it certainly remains possible that TLR2/6, 7/8 or 9 agonists may also have potential for this purpose. Although we postulate Pam3CSK4 and other MyD88-activating TLR agonists exert their adjuvant effects by directly co-stimulating TI-2 Ag-activated B cells, as is the case for MPL, this remains to be formally tested. It is certainly possible that other mechanisms, including induction of IFN known to support these responses (4, 41), are also involved.

Important considerations for developing TLR agonists as vaccine adjuvants to co-stimulate Ag-activated B cells include the effect these agonists have on non-B cell populations and their interaction with other vaccine components—both of which may impact the protective Ab response. Indeed, the unexpected finding that combining TLR2 and TLR7 agonists (ie., adillipoline or R837 plus Pam3CSK4) failed to induce PPS-specific IgG in contrast to when either agonist was used alone highlights the need for careful consideration of TLR agonist combinations. The factors contributing to this antagonism remain unknown. Conversely, Pam3CSK4 exhibited additive effects when combined with the MPL-based adjuvant in mice. Supporting this observation, there are documented instances of positive crosstalk between TLR2 and TLR4 in B cells and other cell types. Notably, TLR2 agonists have been shown to upregulate TLR4 expression on human B cells (42), and the combined stimulation of TLR1/2 and TLR4 has been found to synergistically promote class switch recombination in mouse B cells (43). Indeed, small amounts of contaminating TLR2 and TLR4 agonists in commercial pneumococcal vaccines have been found to support increased Ab responses in mice (44). While the mechanism underlying Pam3CSK4-mediated rescue of MPL-based adjuvant activity in NHP is not yet known, it is possible that in addition to directly co-stimulating Ag-activated B cells via TLR1/2, Pam3CSK4 may have also facilitated MPL activation of primate B cells by upregulating TLR4. Alternatively, Pam3CSK4 may have functioned independently of MPL in the vaccinated NHP. Indeed, the humanized mouse model demonstrated Pam3CSK4-squalene emulsion was sufficient for potent adjuvant effects. While we have only addressed the capacity for boosting of polysaccharide-specific Ab responses in immunocompetent mouse models, future work will investigate the potential for this TLR-based adjuvant to promote long-lived Ab responses and durable, functional memory. Additionally, further work is required to define the mechanisms by which Pam3CSK4 functions in concert with squalene emulsion in the context of polysaccharide vaccines. These investigations are expected to enable optimization of this and other suitable adjuvant alternatives for improving protective humoral immunity to polysaccharide vaccines.

In conclusion, our findings underscore the importance of rational adjuvant design tailored to the unique features of polysaccharide-specific B cell responses. Mechanistic work in mice has been critical for field of adjuvant development. However, key differences in the ability of adjuvants optimized for protein-based vaccines to similarly function as adjuvants for polysaccharide vaccines has hindered progress. By identifying Pam3CSK4 as a potent MyD88-activating adjuvant capable of enhancing IgG responses across species, we provide a compelling foundation for advancing next-generation polysaccharide vaccines. The ability to selectively harness synergistic TLR signaling while avoiding antagonistic combinations opens new avenues for the rapid development of vaccines for emerging pathogens, as well as for providing alternative vaccines, particularly for populations where current conjugate vaccines are limited by availability, cost, complexity, or immunogenicity. Ultimately, our findings lays the groundwork for a more versatile and mechanistically informed approach to eliciting durable, protective humoral immunity against encapsulated bacterial pathogens.

## MATERIALS AND METHODS

### Study Design

Based on our initial experiments focused on evaluating the mechanisms by which MPL-based adjuvants improve polysaccharide-specific Ab responses in mice (via B cell-dependent TLR4/MyD88 activation) and their observed lack of effect in NHP, our goal was to determine whether other MyD88-activating TLR agonists could serve as adjuvants for polysaccharide vaccines in mice, NHP, and humans. For the in vivo studies, we aimed for at least 5 mice per group, based on our previous work with adjuvant evaluation in mice (refs). We followed a similar approach for humanized mice, by using 5 donors for reconstitution studies. AGM were leased with the goal of 4 animals per group due to cost constraints. Animal numbers are included in figure legends. In these studies, biological replicates are considered individual animals, or in the case of humanized mice, individual donor PBMC used to reconstitute an NSG mouse receiving a particular treatment. In vitro assays used one individual donor on different days—each was considered an experimental and biological replicate. The number of times an experiment was repeated is indicated in the figure legend. For mouse experiments, we collected sera out to day 60, based on our previous work. However, we limited serum collection in humanized NSG mice to d28 given the potential for GVHD development. NHP sera were also only collected for primary responses given associated lease costs. Data from animals were included if they were healthy at baseline and throughout the vaccination regimen. Data from one AGM who had an infection and antibiotic treatment during the study was excluded. Data for humanized mice were included if engraftment showed circulating human Ab (total Ab) at day 21.

### Mice

Wild type (WT), mumt, TLR4^-/-^, MyD88^-/-^, Trif^-/-^, and B1-8^hi^ IgH knock-in (B6.129P2-Ptrpca Ightm1Mnz/J) mice on a C57BL/6 background were from the Jackson Laboratory and were bred in-house under SPF conditions. NSG (NOD.Cg-Prkdcscid Il2rgtm1Wjl/SzJ) mice were obtained from the Jackson Laboratory and were maintained on drinking water containing sulfamethoxazole (0.8 mg/ml) and trimethoprim (0.16 mg/ml). Mice were age- and sex-matched for studies. Adult female African Green Monkeys (AGM) were leased from Wake Forest Primate Center’s interventional cohort, as previously described (29). Studies were approved by Wake Forest University School of Medicine’s Animal Use Committee.

### Immunizations and ELISAs

Mice were immunized with the equivalent of ∼0.125 μg each PPS contained within Pneumovax23 (Merck) or 5 μg NP40-Ficoll (Biosearch Technologies) in 50 μl PBS intramuscularly (i.m.) unless otherwise indicated. Sigma Adjuvant System containing 20 μg *Salmonella minnesota* MPL, 20 μg synthetic cord factor (trehalose-6,6‘-dicorynomycolate (TDCM) in 0.5% squalene/0.05% Tween-80 was mixed with Ags prior to injections. Alternatively, adjuvant containing agonist, including Pam3CSK4 (20 μg), R837 (20 μg), adilipoline (20 μg), flagellin, and/or TDB (20 μg) was mixed with 40 μl of the squalene oil in water formulation, Addavax (all from InVivogen). AGM were immunized i.m. (right quadricep) with Pneumovax23 containing ∼12.5 μg each PPS in 250 μl saline (half the human dose) alone or with adjuvant (250 μl Sigma Adjuvant System containing 125 μg MPL and 125 μg TDCM).

Total Pneumovax23- and PPS-specific ELISAs were performed as previously described using cell wall polysaccharide-Multi (SSI Diagnostica) to block cell wall-specific Ab binding (24). NP-specific ELISAs were performed using NP-BSA as a coating antigen as previously described (1, 24). As extensively detailed in our previous publication (24), PPS Ab concentrations were estimated with an indirect method that involves standard curves generated using goat anti-mouse Ig (H+L) or goat anti-human Ig (H+L) capture Abs in conjunction with a dilution series of mouse or human IgM and IgG isotype standards (Southern Biotechnology), respectively. Human Abs were detected using anti-human IgM, IgG, and IgA alkaline phosphatase conjugates (Southern Biotechnology) whereas AGM Abs were detected using anti-monkey IgM, IgG, and IgA Abs (Fitzgerald). Relative PPS-specific IgA levels in AGM were determined using a standard curve developed with a dilution series of positive control sera. Linear, or when more appropriate, polynomial regression was used to estimate the concentration of Abs in test samples.

### Mouse B cell adoptive transfer experiments and bone marrow chimera generation

Mumt mice were reconstituted with purified B cells (EasySep pan-B cell selection kit, StemCell) with 3×10^7^ wild type, TLR4^-/-^, or Trif^-/-^ spleen B cells i.v. and immunized with 5 ug NP_40_-Ficoll i.m. the following day. For bone marrow chimera (BM) generation, WT mice were lethally irradiated (950 rad) and reconstituted i.v. with BM from muMT mice mixed with WT or MyD88^-/-^ BM (80:20 ratio; 1 x 10^7^ BM cells) as previously described (22, 24). Sulfamethoxazole (0.8 mg/ml) and trimethoprim (0.16 mg/ml) was supplied in drinking water 1 week prior to and 2 weeks following irradiation. Mice were rested for 8 weeks prior to immunization.

### Human PBMC and mouse splenocyte cultures/ELISPOTS

Human PBMCs from whole blood or bloody buffy coats (Zenbio, SER-BC-SDS) obtained from healthy donors, aged 21-42, were purified using Lymphoprep (1.077 density). Buffy coats were harvested and washed in PBS. Cells were cultured at 2 to 4 ×10^5^ per 96 well in cRPMI+10% FCS containing 50 μM beta-mercaptoethanol and 10 ng/ml rhuIL-2 (Peprotech). The following agonists were added to cultures, as indicated: Pam3CSK4 (2 μg/ml), FSL-1(1 μg/ml), flagellin (1 μg/ml), R837 (1 μg/ml), R848 (1 μg/ml), CpG2006 (2 μg/ml), Adilipoline (1 μg/ml), Addavax (1:500, equivalent to 0.01% squalene), biotinylated goat anti-human Ig (H+L) F(ab’)_2_ (1 μg/ml; Jackson Immunoresearch), and/or streptavidin (5 μg/ml). In some experiments, PBMC were labeled with CFDA, SE (CFSE, 1 μM; Invitrogen) prior to culture. B cells were purified from PBMC using Miltenyi’s CD19 positive selection kit according to manufacturer’s instructions. Cultures were incubated for 5 days and cells were harvested for flow cytometric analysis (described below) or further culture in ELISPOT plates. For ELISPOTs, 20 μl cells were harvested on day 5 and placed in a PVDF Immobulon P plate (MSIPS4510) coated with goat anti-human Ig (H+L) (5 μg/ml) and preblocked with 180 μl media. Plates were incubated for 18 hours in a CO_2_ incubator at 37C and developed using anti-human IgM, IgG, and IgA alkaline phosphatase conjugates (Southern Biotechnology), followed by NBT/BCIP development. For mouse B cell cultures, splenocytes were isolated from V_H_B1-8 Tg mice and cultured alone or with NP-Ficoll (10 ng/ml), TLR agonists, or both for 5 days and NP-specific IgM and IgG was measured in supernatants by ELISA as previously described (24).

### Flow cytometry

PBMC cultures were harvested in PBS containing 2% newborn calf serum with the addition of Countbright beads, incubated with normal mouse serum for 15 minutes, and stained with fluorochrome-conjugated Abs specific for CD20 (2H7), CD86 (Fun1), IgG (G18-145), IgM (MHM-88), (CD11b (M1/70) as well as Live/Dead Aqua for 30 minutes at 4C. Fluorochrome-labeled isotype controls were used to determine background staining levels. Cells were analyzed using a FortessaX20 cytometer (BD Biosciences) with FSC-A/FSC-H doublet exclusion. Data were analyzed using FlowJo analysis software (Tree Star).

### huPBMC NSG chimeras

NSG mice were injected with ∼10^7^ freshly isolated donor huPBMC i.p. Mice were immunized the following day with Pneumovax23, Pam3CSK4 + Addavax, or both i.p., with sera collected at 7 day intervals.

### *Streptococcus pneumoniae* challenge

Mice were infected i.n. with serotype 3 WU2 strain *S. pneumoniae* by distributing 40 μl bacteria (1 x 10^7^ CFU) between 2 nares of isofluorane-anaesthetized mice (22). Systemic pneumococcal infections and passive serum protection experiments were carried out by transferring sera and 1000 CFU TIGR4 or 100 CFU WU2 *S. pneumoniae* i.p. (100 μl) into CD19^-/-^ mice as previously described (23). Mice were monitored daily for signs of distress and humanely euthanized.

### Statistical analyses

Data are shown as means ± SEM, with individual data points shown in bar graphs. Differences between sample means were assessed using Student’s t test, Mann Whitney or one-or two-way ANOVA with post hoc analysis as indicated in the figure legends. Differences in survival was assessed using the Log Rank test. Statistical analysis was performed using GraphPad Prism (version 10).

## List of Supplementary Materials

Figures S1-S3 data files.

## Author contributions

K.M.H. conceptualized the study, acquired funding, performed the investigation, supervised the project, visualized the data, and wrote the original draft of the manuscript. JEJ-G and AEA-S performed the investigation, visualized the data, and reviewed and edited the manuscript.

## Data availability

All data associated with this study are present in the paper or the Supplementary Materials.

