## Supplemental Figures 1-3 for "Pam3CSK4 as a Cross-Species Adjuvant for Polysaccharide Vaccines: Efficacy in Humanized Mouse and Non-Human Primate Models"

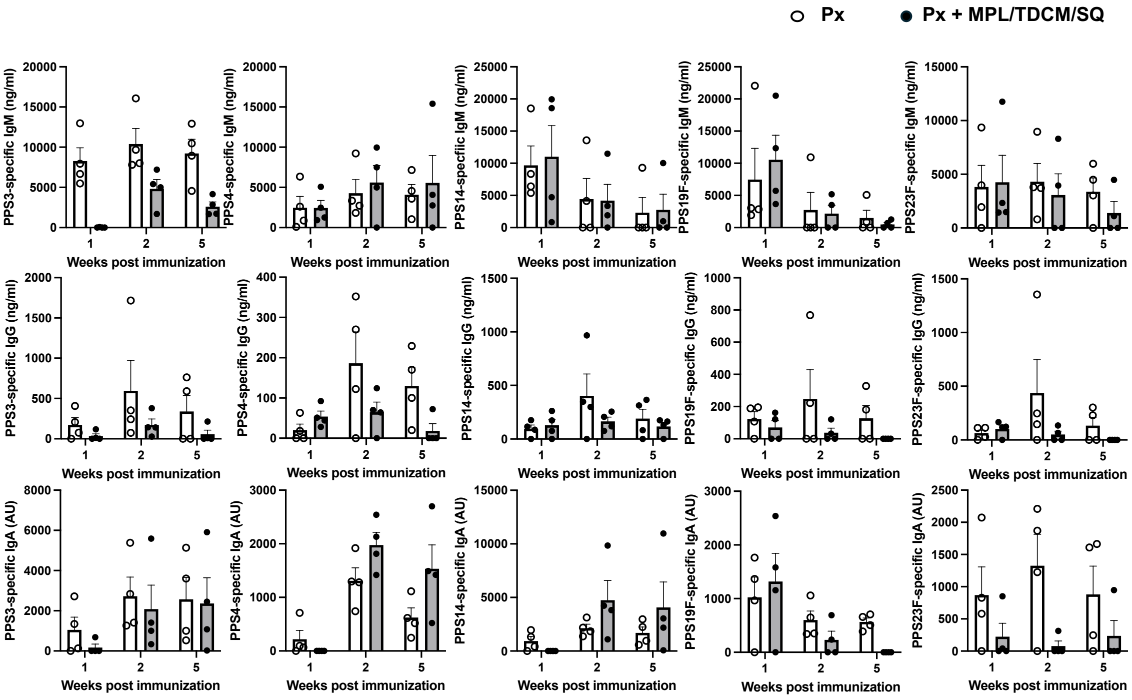


**Supplemental Figure 1. MPL+ TDCM + squalene emulsion does not increase PPS3-, 4-, 14, 19F, or 23F-specific Ab responses in African Green monkeys (AGM).**
PPS-specific IgM, IgG, and IgA responses in 17-20 year old female AGM 1, 2, and 5 weeks post-vaccination (i.m.) with Pneumovax23 containing 12.5 µg each PPS alone or mixed with MPL+TDCM+squalene emulsion (SQ) adjuvant. Individual baseline Ab values were subtracted from week 1, 2, and 5 Ab values to determine increases in PPS-specific IgM, IgG, and IgA levels over baseline as shown in graphs (n=4 AGM/group). Circles indicate values for individual animals.

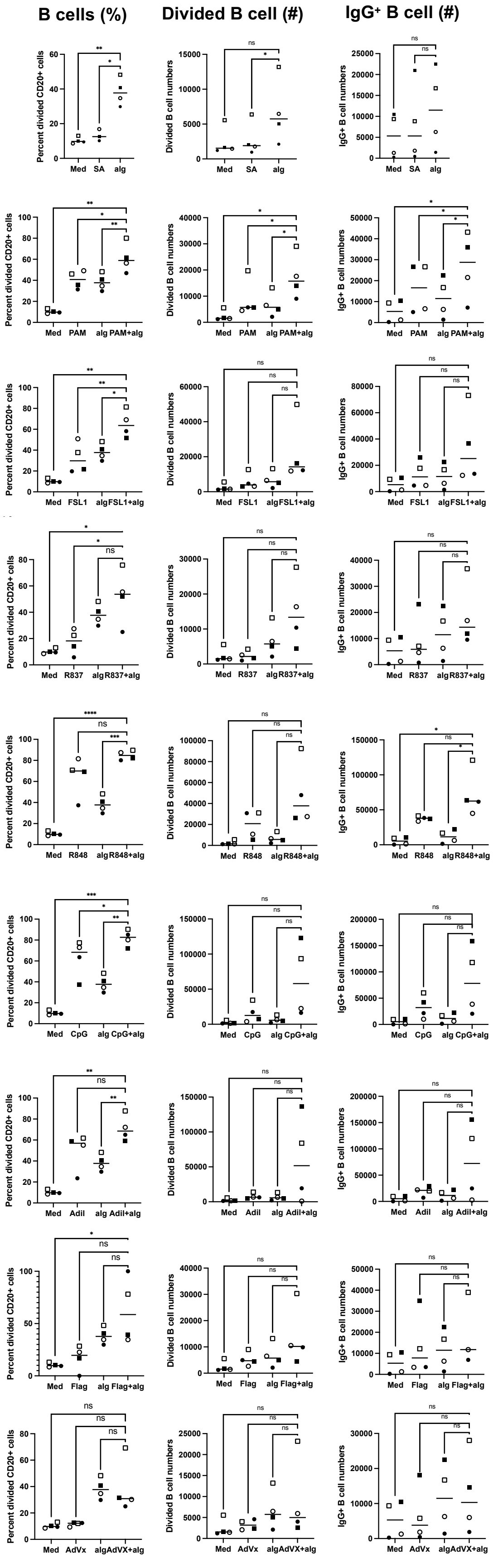


**Supplemental Figure 2. Expansion of human B cells in PBMC with TLR agonists, biotinylated anti-Ig (H+L) + streptavidin, or both for 5 days.** PBMC from 4 human donors were labeled with 1 uM CFSE and cultured alone with rhuIL-2. TLR agonists, biotinylated anti-Ig (H+L) + streptavidin, or both were added for 5 days. Cells were harvested for flow cytometric analysis, and analyzed for viable $CD20^+$ B cells, CFSE-divided B cell and $IgG^+$ cell numbers. Data were analyzed by one way ANOVA with comparisons draw for TLR agonist + BCR-activated B cells versus other conditions. Asterisks indicate significant differences (*p<0.05, **p<0.01, ***p<0.001).

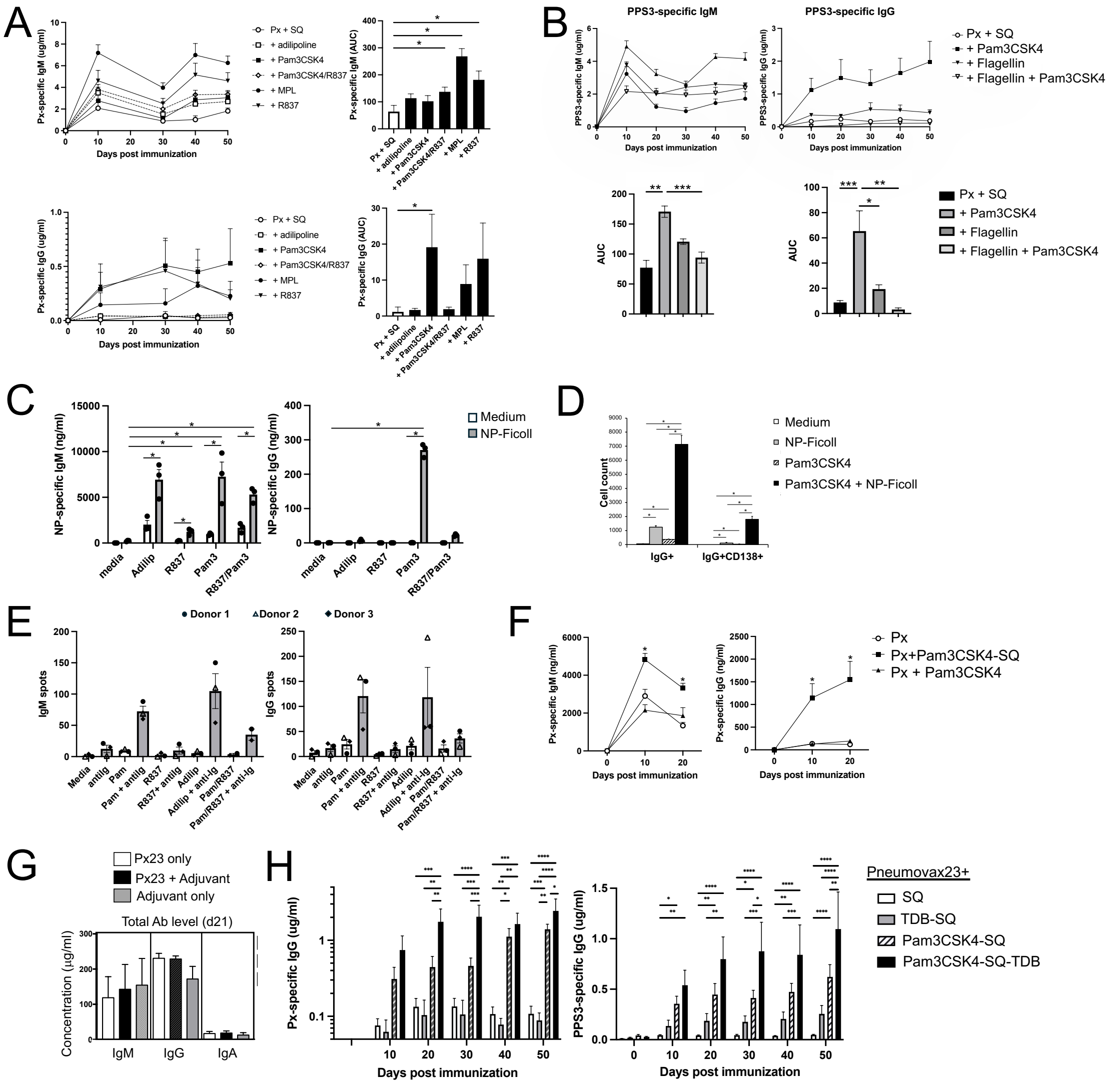


**Supplemental Figure 3. Effects of adjuvants on PPS-specific TI-2 Ab responses.**
A-B) Pneumovax23(Px) (A) or PPS3 (B) -specific IgM and IgG responses in WT mice immunized i.m. with Px (0.125 µg each PPS) plus squalene emulsion (SQ) emulsion (40 µl) alone or combined with the indicated TLR agonists. Differences in responses between mice receiving TLR agonists versus mice immunized with Px+SQ alone were assessed by area under the curve (AUC) analysis and one-way ANOVA with Bonferroni's post-hoc analysis (n=4-6 mice/group). C-D) $V_H$B1-8 Tg splenic B cells were cultured with NP-Ficoll, adilipoline, Pam3CSK4, R837, Pam3CSK4 plus R837 individually or combined for 5 days. NP-specific IgM and IgG production was assessed in supernatants by ELISA (C) and NP-specific B cells were assessed for class switching and differentiation to ASC ($CD138^+$) by flow cytometry (D). E) Pam3CSK4-supported increases in anti-Ig-induced ASC in ELISPOT analysis in purified human B cell cultures (d6) were reduced when R837 was added. In C and E, circles indicate individual mice or human donors (n=3/group). F) PPS3-specific IgM and IgG responses in WT mice in response to Px, Px+Pam3CSK4, Px+Pam3CSK4+SQ. G) Circulating human Ab levels (mean + SEM) in NSG mice 3 weeks post reconstitution (n=5 donors). H) Px-and PPS3-specific specific IgG levels in response to Px mixed with different adjuvants. Asterisks indicate significant differences as assessed by repeated measures ANOVA with post-hoc analysis. Asterisks indicate significant differences (*p<0.05, **p<0.01, ***p<0.001,****p<0.0001).
